# Automatic quantification of disgust reactions in mice using machine learning

**DOI:** 10.1101/2023.04.23.538002

**Authors:** Shizuki Inaba, Naofumi Uesaka, Daisuke H. Tanaka

**Affiliations:** Department of Cognitive Neurobiology, Graduate School of Medical and Dental Sciences, Institute of Science Tokyo, 1-5-45 Yushima, Bunkyo-ku, Tokyo 113-8519 Japan

## Abstract

Disgust, a primary negative emotion, plays a vital role in protecting organisms from intoxication and infection. In rodents, this emotion has been quantified by measuring the specific reactions elicited by exposure to unpleasant tastes. These reactions were captured on video and manually analyzed, a process that required considerable time and effort. Here we developed a method to automatically count disgust reactions in mice by using machine learning. The disgust reactions were automatically tracked using DeepLabCut as the coordinates of the nose and both front and rear paws. The automated tracking data were split into test and training data, and the latter were combined with manually labeled data on whether a disgust reaction was present and, if so, which type of disgust reaction was present. Then, a random forest classifier was constructed, and the performance of the classifier was evaluated in the test dataset. The total number of disgust reactions estimated by the classifier highly correlated with those counted manually (Pearson’s r = 0.98). The present method will decrease the time and effort required to analyze disgust reactions, thus facilitating the implementation of the taste reactivity test in large-scale screening and long-term experiments that necessitate quantifying a substantial number of disgust reactions.

## Introduction

Disgust, a primary negative emotion recognized since Darwin [1], is elicited by a distaste for food and contact with feces. This emotion plays a crucial role in protecting animals from intoxication and infection by helping them avoid harmful substances and pathogens [2]. Abnormal manifestations of disgust have been observed in patients receiving chemotherapy for cancer [3–4] and have been proposed in the pathogenesis of various intractable psychiatric disorders, including eating disorders and obsessive-compulsive disorder [5–7]. Therefore, elucidating the neural mechanisms underlying disgust is essential for understanding the biological basis of emotion, reducing drug side effects, and effectively managing various psychiatric disorders.

Since disgust is a conscious experience and thus cannot be directly observed [8–12], an observable readout is required to elucidate its neural mechanisms. Taste reactivity, consisting of orofacial and somatic behavioral reactions in response to taste stimuli, expresses the hedonic evaluation of taste, but not taste quality [13–15]. Several specific behaviors, such as gape, in taste reactivity have been established as reliable indicators of disgust [15] and are referred to as disgust reactions [16–19].

The quantification of disgust reactions is so complex that manual analysis has been carried out and automated analysis methods have not been developed. In previous analyses of disgust reactions, orofacial and somatic behavioral reactions were video-recorded, and the relevant behaviors were subjectively distinguished while the video was played frame-by-frame. Thus, the quantified data of a particular test can be somewhat variable between the people analyzed, and a significant amount of time and effort is required for the analysis [20].

To overcome these limitations and reproducibly and quickly assess disgust reactions in mice, we constructed a classifier to automatically count them. To accomplish this, we utilized DeepLabCut as a pose estimation system based on transfer learning with deep neural networks that track defined body parts of animals without physical markers [21] and random forest as a supervised learning model with an ensemble learning method [22].

## Methods

### Animals

Adult (6-7 weeks-old) male C57BL/6J mice (n = 27) were obtained from Japan SLC Inc. (Shizuoka, Japan). Upon arrival, mice were housed in clear plastic cages (18 × 26 × 13 cm) with wood tips (Soft tip: Japan SLC, Shizuoka, Japan) in groups of 2-4 males per cage and were then housed individually in smaller cages (14 × 21 × 12 cm) immediately before handling. Mice were maintained at 23 ± 1 °C under a 12 h light/dark cycles (lights on at 8:00 am) and given *ad libitum* access to food (Labo MR stock; Nosan Corp., Kanagawa, Japan) and tap water. The tap water was changed to Milli-Q water 3 days before beginning the reaction experiments. All behavioral experiments were performed during the light cycle. All the animal experiments were approved (No. 0150384A, 0160057C2, 0170163C, A2017-194A, and A2018-138C4) by the Institutional Animal Care and Use Committee of the Institute of Science Tokyo, and was performed according to the ARRIVE and relevant guidelines and regulations.

### Surgery for intraoral tubing

For implantation of an intraoral tube, mice were anesthetized by intraperitoneal (i.p.) injection of sodium pentobarbital (40 mg/kg BW; Nembutal; Abbott Laboratories) or a mixture of midazolam (4 mg/kg BW; Astellas Pharma Inc.), butorphanol (5 mg/kg BW; Meiji Seika Pharma Co., Ltd.), and medetomidine (0.3 mg/kg BW; Nippon Zenyaku Kogyo Co., Ltd.) [23]. The depth of anesthesia was maintained at a sufficient level to prevent paw pinch reflex. An incision was made in the midline of the scalp. A curved needle attached to an intraoral polyethylene tube (SP-10; Natsume Seisakusho Co., Ltd., Tokyo, Japan) was inserted from the incision site and advanced subcutaneously posterior to the eye, to exit at a point lateral to the first maxillary molar on the right side of the mouth. The intraoral end of the tube was heat-flared to an approximate diameter of 1 mm to prevent it from drawing into the oral mucosa. Mice were mounted in a stereotaxic frame (David Kopf Instruments, Tujunga, CA, USA) using ear bars. Two alcohol-sterilized small screws (PN-04; LMS Co., Ltd., Tokyo, Japan) were anchored to the skull and a piece of plastic bar was fixed to the screws with dental cement (Unifast3; GC Corp., Tokyo, Japan). The end of the tube exiting the head incision was fixed to a plastic bar with dental cement. Mice received subcutaneous injections of the antibiotic chloramphenicol sodium succinate (60 mg/kg BW, Chloromycetin Succinate; Daiichi Sankyo Co., Ltd., Tokyo, Japan) for infection prevention and carprofen (5 mg/kg BW, Rimadyl; Zoetis, Tokyo, Japan) for pain relief. Three days after surgery, one end of a delivery tube (SP-10; Natsume Seisakusho Co., Ltd.) was connected to the end of the intraoral tube fixed to the plastic bar on the mouse head using a connector (KN-394, two directions (0.3+0.3); Natsume Seisakusho Co., Ltd.), and the other end was connected to a needle (30Gx13mm; ReactSystem Co., Osaka, Japan) attached to a 1-mL syringe, and ∼20 µL of sterile MilliQ water was introduced into the mouth of the animal to test the patency of the tubes. The intraoral tube was flushed with ∼20 µL sterile MilliQ water every 2-3 days to prevent occlusion. The mice were allowed 1-3 weeks to recover from surgery before the beginning of the behavioral experiments.

### Taste reactivity test

The taste reactivity test [14,15,18] was used to measure affective behavioral reactions in mice. The test chamber was composed of a glass floor and an acrylic cylinder (30 cm height, 10 cm outside diameter, and 3 mm thickness). A digital video camera (HDR-PJ800; Sony, Tokyo, Japan) was placed beneath the glass floor to record the ventral view of the mouse. One end of the delivery tube was connected to the end of the intraoral tube fixed to the plastic bar on the mouse head using a connector, and the other end was connected to a needle attached to a 1-mL syringe. The stimulant solution was filled into a syringe and infused into the mouth of the mice. Orofacial and somatic behaviors during the infusions were video-recorded in a bottom-up view at 30 frames per second (Fig. 1A).

**Figure 1.**
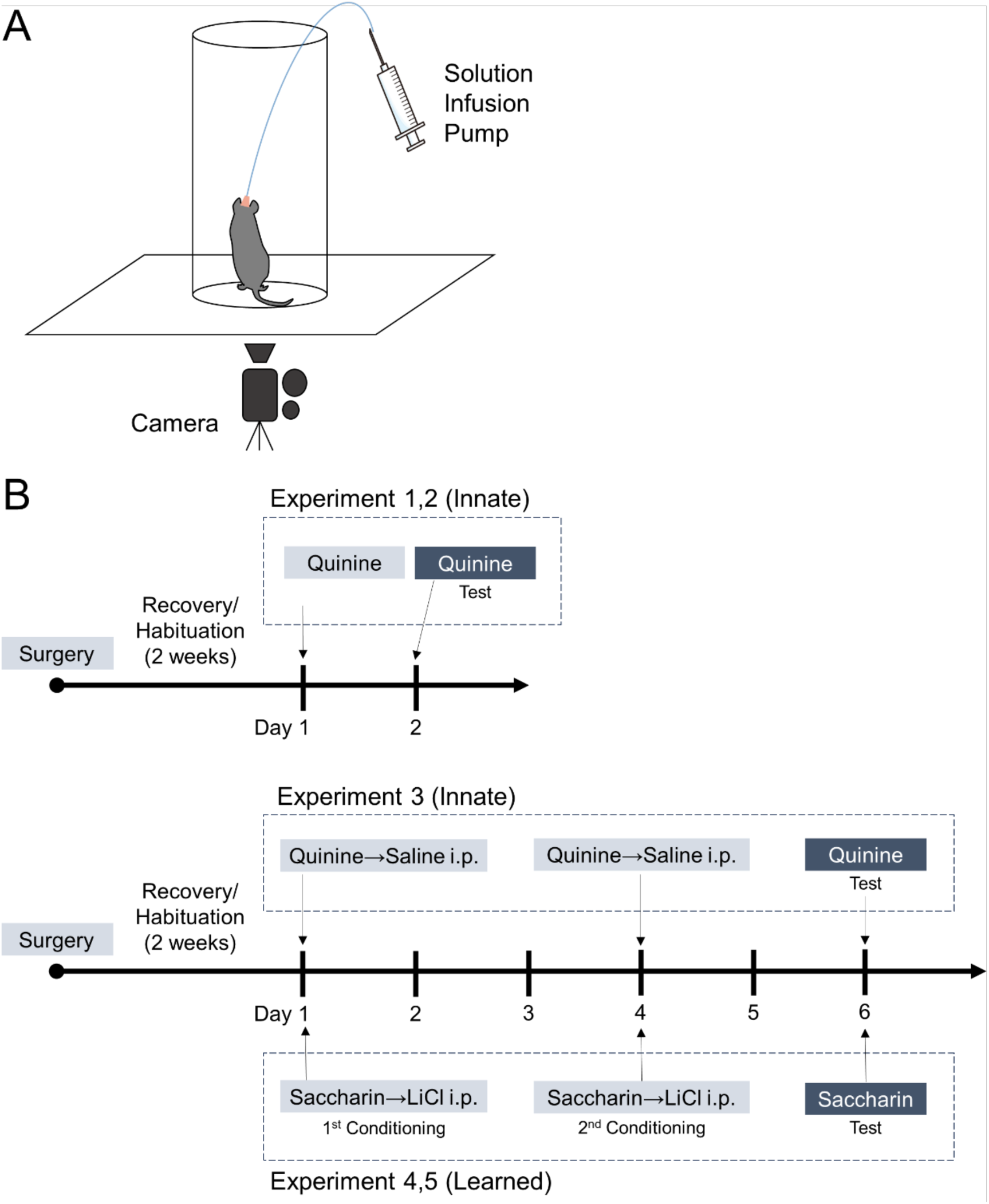
Experimental setup for taste reactivity test in mice. **(A)** Solution was infused intraorally through an implanted tube. Orofacial and somatic behaviors during infusions were video-recorded in a bottom-up view. **(B)** This scheme shows the Experimental timeline. <u>Experiment 1,2 (Innate)</u>: On Day 1, mice received intraoral infusions of Quinine solution as habituation. On Day 2 (test day), the mice received the same infusions as those performed on Day 1, and the behaviors of the mice during the infusions were video-recorded. In Experiment 1, infusions were performed twice for a total of 1 min each, with a 1 min interval in between. In Experiment 2, the infusions were continuous for 2 min, without intervals. <u>Experiment 3 (Innate)</u>: On Day 1, mice received intraoral infusions of quinine solution, and shortly thereafter received an i.p. injection of physiological saline as a control treatment for the learned groups described below. The same procedure was repeated on Day 4. On Day 6 (test day), the mice received intraoral infusions of quinine solution, and their behaviors during the infusions were video-recorded. In Experiment 3, infusions were performed twice for a total of 1 min each, with a 1 min interval in between. <u>Experiment 4,5 (Learned)</u>: On Day 1, mice received intraoral infusions of saccharin solution, and shortly after that, they received an i.p. injection of LiCl solution (1^st^ conditioning). The same procedure was repeated on Day 4 (2^nd^ conditioning). On Day 6 (test day), the mice received intraoral infusions of saccharin solution, and their behaviors during the infusions were video-recorded. In Experiment 4, infusions were performed twice for a total of 1 min each, with a 1 min interval in between. In Experiment 5, the infusions were continuous for 2 min without intervals.

To assess disgust reactions in the innate and learned experimental conditions, the mice were divided into an innate group (n = 13) and a learned group (n = 14) after surgery. As the experiments were conducted while optimizing the experimental conditions, the conditions differed from experiment to experiment. Three different schedules and experiments were performed for the innate groups and two slightly different schedules and experiments were used for the learned groups (Fig. 1B, Table 1). Each mouse was used in a single experiment. The number of mice used in each experiment was determined by unintentional factors such as the number of mice available at the time the experiment was performed. The experimental schedule for each experiment is as follows.

**Table 1.** Taste reactivity test schedule.

|  | Innate |  | Learned |
| --- | --- | --- | --- |
|  | Expt. 1,2 | Expt. 3 | Expt. 4,5 |
|  | Surgery |  |  |
|  | Recovery / Habituation |  |  |
| Day 1 | Q | Q +Saline | S +LiCl |
| Day 2 | Q (Test) |  |  |
| Day 3 |  |  |  |
| Day 4 |  | Q +Saline | S +LiCl |
| Day 5 |  |  |  |
| Day 6 |  | Q (Test) | S (Test) |

This table presents an experimental timeline. All mice underwent the same surgeries, recoveries, and habituations. On the final experimental (Test) day, all mouse behaviors were video-recorded for analysis.
Q: Intraoral quinine infusion, S: Intraoral saccharin infusion, +Saline: Saline i.p. immediately after intraoral infusion, +LiCl: LiCl i.p. immediately after intraoral infusion.

#### Experiment 1 for the innate group

On Day 1, mice (n = 5) received intraoral infusions of 50 µL of 3 mM quinine solution twice for 1 min each with a 1 min interval in between to adapt to the test procedure. On Day 2 (test day), the mice received the same infusions as those administered on Day 1, and their behavior during the infusions was recorded (Fig. 1B, Table 1).

#### Experiment 2 for the innate group

On Day 1, mice (n = 2) continuously received infusions of 80 µL of 3 mM quinine solution for 2 min without an interval to adapt to the test procedure. On Day 2 (test day), the mice received the same infusions as those administered on Day 1, and their behaviors during the infusions were video-recorded.

#### Experiment 3 for the innate group

On Day 1, mice (n = 6) received intraoral infusions of 50 µL of 3 mM quinine solution twice for 1 min each with a 1 min interval in between. Shortly thereafter, the mice received an intraperitoneal (i.p.) injection of physiological saline (10 mL/kg body weight [BW]) as a control treatment in the learned group. The same procedure was repeated on Day 4. On Day 6 (test day), the mice received the same infusions as those administered on Day 1, and their behaviors during the infusions were video-recorded.

#### Experiment 4 for the learned group

On Day 1, mice (n = 11) received infusions of 50 µL of 5.4 mM saccharin solution twice for 1 min each with a 1 min interval in between. Shortly thereafter, the mice received an i.p. injection of LiCl solution (0.3 M, 10 mL/kg BW) (1^st^ conditioning). The same procedure was repeated on Day 4 (2^nd^ conditioning). On Day 6 (test day), the mice received the same infusions as those administered on Day 1, and their behaviors during the infusions were video-recorded.

#### Experiment 5 for the learned group

On Day 1, mice (n = 3) continuously received infusions of 80 µL of 5.4 mM saccharin solution for 2 min without an interval. Shortly thereafter, the mice received an i.p. injection of LiCl solution (0.3 M, 10 mL/kg BW) (1^st^ conditioning). The same procedure was repeated on Day 4 (2^nd^ conditioning). On Day 6 (test day), the mice received the same infusions as those administered on Day 1, and their behaviors during the infusions were video-recorded.

Thus, videos obtained from five different experiments were used for subsequent analysis. Among these videos, almost half (n = 14; innate = 7, learned = 7) were obtained in a previous study in which automated tracking was not performed [18] (Table 2).

**Table 2.** Videos used in the present studies.

| Video title | Experiment | Innate or learned | Training or Test | Published or unpublished |
| --- | --- | --- | --- | --- |
| 299 | 1 | Innate | Train | Published |
| 309 | 1 | Innate | Train | Published |
| 332 | 1 | Innate | **Test** | Published |
| 334 | 1 | Innate | **Test** | Published |
| 343 | 1 | Innate | Train | Published |
| 529 | 2 | Innate | **Test** | Published |
| 534 | 2 | Innate | Train | Published |
| 251 | 3 | Innate | Train | Unpublished |
| 264 | 3 | Innate | Train | Unpublished |
| 281 | 3 | Innate | **Test** | Unpublished |
| 363 | 3 | Innate | Train | Unpublished |
| 366 | 3 | Innate | Train | Unpublished |
| 367 | 3 | Innate | Train | Unpublished |
| 246 | 4 | Learned | Train | Unpublished |
| 247 | 4 | Learned | Train | Unpublished |
| 250 | 4 | Learned | **Test** | Unpublished |
| 263 | 4 | Learned | Train | Unpublished |
| 278 | 4 | Learned | Train | Unpublished |
| 280 | 4 | Learned | Train | Unpublished |
| 365 | 4 | Learned | Train | Unpublished |
| 293 | 4 | Learned | Train | Published |
| 303 | 4 | Learned | **Test** | Published |
| 324 | 4 | Learned | **Test** | Published |
| 326 | 4 | Learned | Train | Published |
| 406 | 5 | Learned | Train | Published |
| 448 | 5 | Learned | **Test** | Published |
| 450 | 5 | Learned | Train | Published |

Published, these videos were recorded by Tanaka et al. (2019), in which automated tracking was not performed.

### Automated body part tracking

DeepLabCut2.2rc1 (DLC) [23] was used to develop a tracking framework with five mouse-body parts (Fig. 2A). Frame choice and training were performed using the default settings of the DLC. Initially, 20 frames were selected from each video for training purposes. The provisional coordinates of the mouse body part were estimated according to the temporarily created model, and 40 candidate frames for false predictions were selected for each video and manually annotated. This procedure was repeated until the loss was <0.001 and the error was sufficiently small. After the training sessions were completed, the coordinates of the mouse body part in the other (not manually annotated) frames were estimated (Fig. 2A).

**Figure 2.**
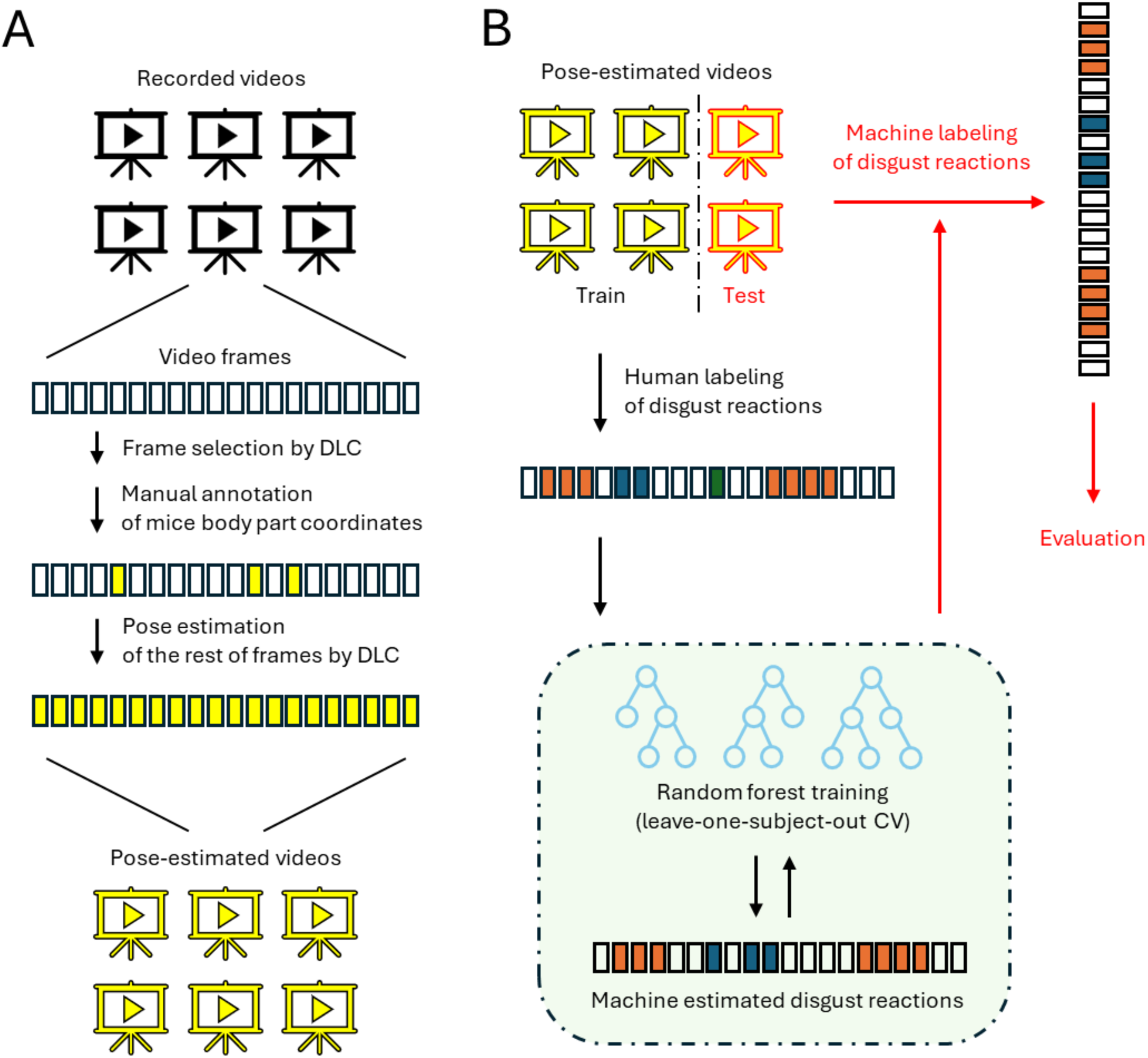
Schematic diagram of the machine learning estimation method used. **(A)** Schematic illustration of pose estimation using DLC. **(B)** Schematic illustration of the Disgust Classification of each frame using random forest.

### Disgust reaction labeling

All 27 videos, including those taken in a previous study [18], were (re)analyzed by a blinded observer who was not involved in the previous study [18]. According to previously established rules [15,18], all video frames were labeled manually frame-by-frame (30 frames/sec) to determine whether they constituted disgust reactions, and if so, which of the five types of disgust reactions they were. The five types of disgust reactions were gapes (large opening of the mouth with retraction of the lower lip), headshakes (rapid lateral movement of the head), face washes (wipes over the face with the paws), forelimb flails (rapid waving of both forelimbs), and chin rubs (pushing the chin against the floor of the test chamber).

### Random forest classification and model evaluation

We extracted 25 per-frame geometric features. They consisted of five displacements of each body part (i.e., nose, left front paw, right front paw, left rear paw, and right rear paw) from the previous frame, ten distances between each body part (e.g., the distance between the nose and left front paw), and ten amounts of changes of each distance from the previous frame. In addition, for each frame, the sum of all features from five consecutive frames, including the two frames before and after the frame, was considered, so that each frame had 125 features.

We then constructed a random forest classifier using the caret [24] and Rborist [25] packages in R. Before developing the classifier, the data were randomly divided into training (n = 19; innate = 9, learned = 10) and test (n = 8; innate = 4, learned = 4) datasets, with an almost equal number of the innate and learned groups (Table 2). The classifier was constructed by setting the classWeight argument in the train function of the caret package to ‘balance’ in view of the imbalanced presence of frames containing movements of interest, which allowed the classes to be weighted inversely proportional to their frequency of occurrence in the data. In the training sessions, the dataset was leave-one-subject-out (19-fold) cross-validated for each mouse to prevent overfitting (Fig. 2B). In addition, we performed hyperparameter tuning and selected the model with the highest accuracy as the final one using the tuneLength argument in the train function of the caret package. Although the number of ensembled decision trees was fixed at the default value of 500 during the model creation process, we confirmed that the number of ensembled decision trees was sufficient by separately training after the final model creation, with the same training parameters changing only nTree.

After developing the final model, we evaluated the classifier’s frame-by-frame performance in discriminating each type of disgust reaction by calculating the overall accuracy, as well as its positive predictive value (PPV) and sensitivity. PPV, also called precision, is expressed by the following equation:

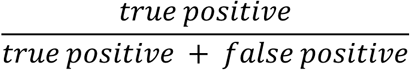

Sensitivity, also called recall, is expressed by the following equation:

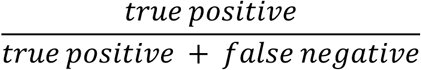

### Disgust reaction scoring

For the five types of disgust reactions, we analyzed consecutive frames with the same label as one count. To ensure that each component of disgust reactions contributed equally to the final scores, reactions that occurred in continuous counts were scored in time bins according to previously established rules [15,26]. Components characterized by long-duration counts, such as chin rubs, were scored in five-second bins (successive repetitions within five seconds scored as one occurrence), and face wash was scored in two-second bins. The other three reactions that could occur as a single behavior were scored as separate occurrences (e.g., one forelimb flail equals one occurrence). The total disgust reaction scores were quantified as the sum of all five binned counts during two minutes of intraoral infusion of the test solutions.

### Computer software and hardware

A desktop computer equipped with an Intel Core i7-10700 CPU, an NVIDIA GeForce GTX 1660 SUPER GPU, and 16 GB of RAM was used for mouse posture estimation in DLC and data processing in R.

## Results

### Manual analysis of disgust reactions

First, we manually and quantitatively analyzed 97200 frames from 27 videos that recorded disgust reactions. Consistent with a previous study [18], disgust reactions were observed in both the innate and learned groups with a similar number of reactions for each type of disgust reaction (Fig. 3A). Regarding the number of counts for each type of disgust reaction, forelimb flails, face washes, and chin rubs were more common than gapes and head shakes (Fig. 3B, Wilcoxon signed-rank sum test; gape vs. chin rub: p=0.049, gape vs. face wash: p<1e-6, gape vs. forelimb flail: p<1e-7, head shake vs. chin rub: p=0.031, head shake vs. face wash: p<1e-7, head shake vs. forelimb flail: p<1e-7).

**Figure 3.**
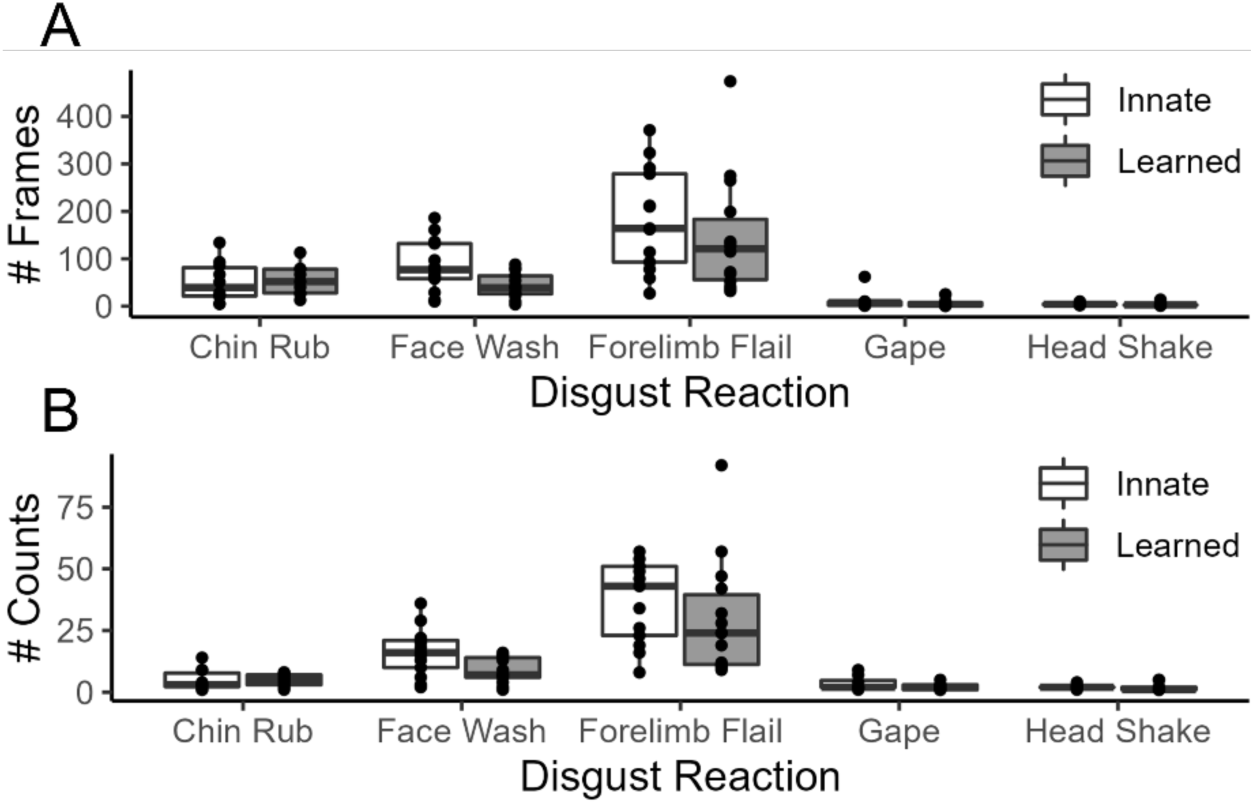
The number of each type of disgust reaction. **(A)** Number of frames that included any disgust reactions confirmed manually. **(B)** The number of counts for each type of disgust reaction was confirmed manually. Each dot represents a value for a mouse.

### Automated body part tracking

Next, we automatically estimated the mouse posture to perform automatic quantification of video-recorded disgust reactions. Because the optimal number of body parts to be labeled for posture estimation is yet to be well established, we referred to the occurrence rate of each type of disgust reaction (Fig. 3B) to determine the number of markers. The movements that can be tracked by focusing on the nose and the front and rear paws (i.e., chin rubs, face washes, forelimb flails, and head shakes) comprised 96.23% of the disgust reactions. In contrast, movements that required attention to mouth movements (i.e., gapes) were rarely observed (3.77%). Thus, we defined the five body parts of mice, the nose, and the front and rear paws (Fig. 4A) were sufficient for the objective of the present study, which estimated the total scores for the disgust reaction.

**Figure 4.**
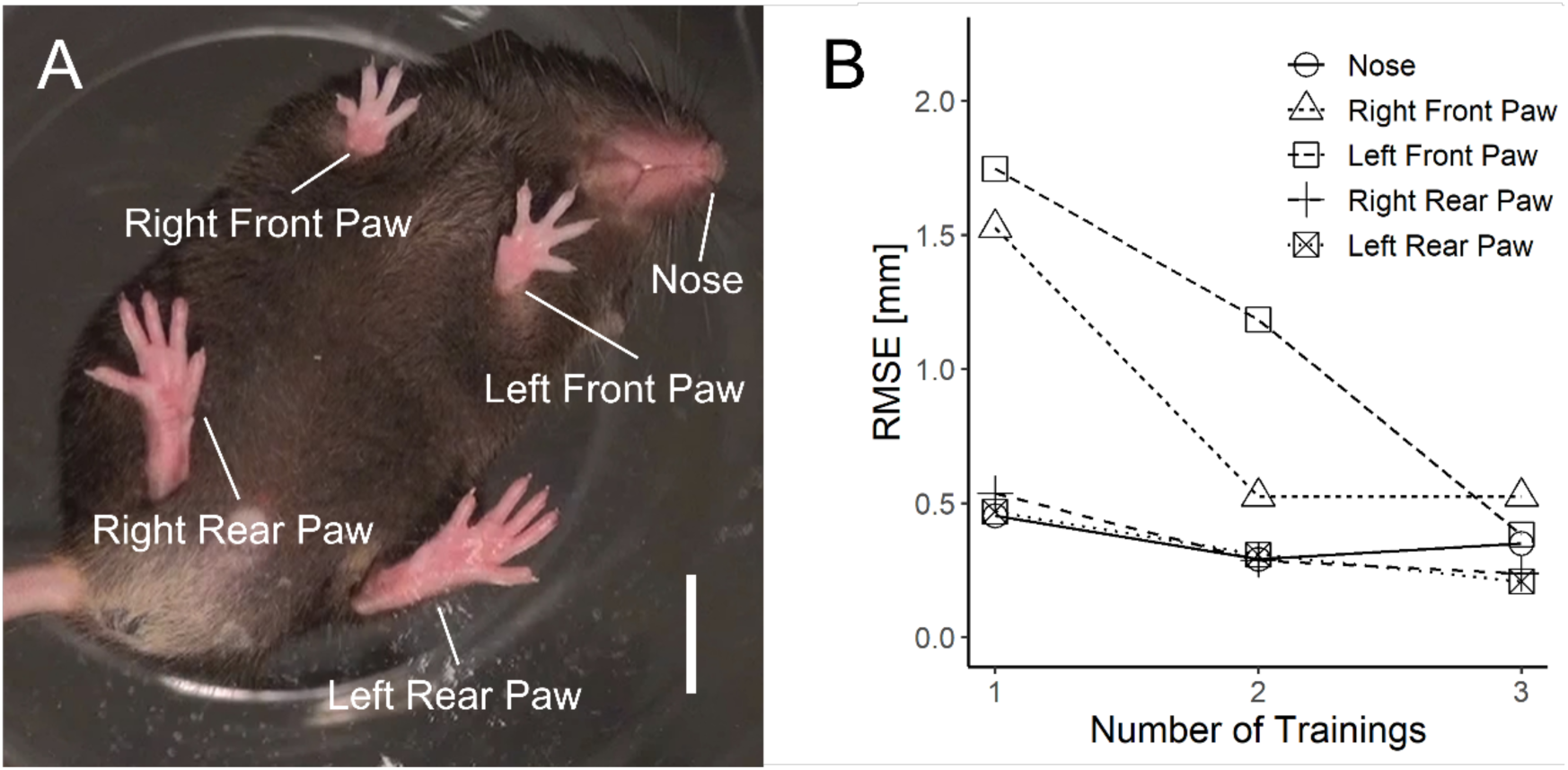
Annotation of five body parts for pose estimation. **(A)** Definition of body parts used to estimate mouse posture using the DLC. Only those visible in the bottom-up view were annotated. Scale bar, 1 cm. **(B)** Root mean square error (RMSE) for each body part during the training.

In the DLC training process, 2674 frames from 27 single-trial video recordings were manually annotated through three training sessions with 500000 iterations each (Fig. 2). The pose-estimation errors were evaluated after the model training procedure. The results showed that the mean error for each body part was less than 1 mm (Fig. 4B), which was sufficiently small compared to previous studies using DeepLabCut (1-8 mm) [27–28] to confirm that the training was complete.

### Outlier frame detection and correction

Next, we corrected for mouse posture estimation because our posture estimation often failed in specific postures (e.g., standing up with its paws against a wall) and rapid movements that could not be captured on a recording. We introduced a rule-based correction procedure that refers to the likelihood of preventing incorrect estimations in such cases. Specifically, the following two corrections were made for all frames in the video posture-estimated by DLC when the estimated likelihood of a particular body part in a specific frame was less than a predetermined threshold value. (1) If the likelihood of the body part in the next frame exceeded the threshold value, the position of the body part in the frame was corrected in the middle of the frame before and after. (2) If the likelihood of the body part in the next frame was below the threshold, the body part’s position in the frame was corrected to the same position as that in the previous frame. In the present study, we set the threshold value to 0.6, which is the default value for an outlier correction method inside the DLC.

### Random forest classifier

We developed a random forest classifier to automate the image classification task of determining whether each video frame contained the behaviors of interest. Before developing the classifier, the data were randomly and equally divided into training and test datasets between the innate and learned groups. We developed three random forest models with different predFixed hyperparameters, that is, the number of randomly selected features when forming each split in a classification tree, of the Rborist package in R. Finally, we adopted a model with a parameter value of 63 and the highest cross-validated accuracy (Fig. 5). To confirm that 500 was sufficient for the number of decision trees ensembled in the final model, separate training was conducted with nTree varying from 10 to 1000 in steps. The results confirmed that there was no significant difference in Accuracy for nTree > 200.

**Figure 5.**
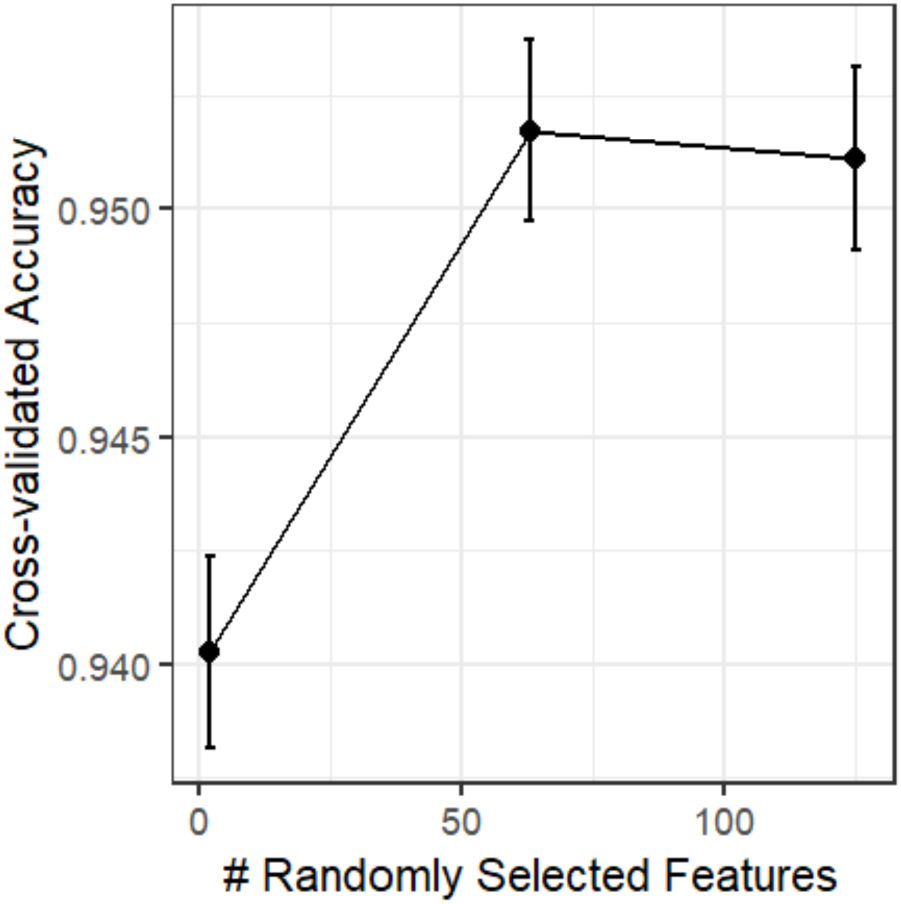
The relationship between the number of randomly selected features and cross-validated accuracy for hyperparameter tuning. The dots represent the mean accuracy of the entire cross-validation and the vertical lines represent the standard deviation.

After the random forest model was completed, the importance of the features was checked using the varImp function in the Carete package to find the features that are likely to contribute strongly to the performance of this model. The importance of each feature is determined from the degree of degradation in the prediction performance when only that feature is randomly shuffled from the original data. This is based on the hypothesis that if the feature is very useful, the prediction using the data containing the shuffled feature should have a much worse performance than the prediction using the original data that was not shuffled. The most important features were the distance between the two front paws and the distance between the nose and right/left front paw (Fig. 6), indicating that the positional relationship between the two front paws and the nose of the mouse was particularly important for the prediction of this model.

**Figure 6.**
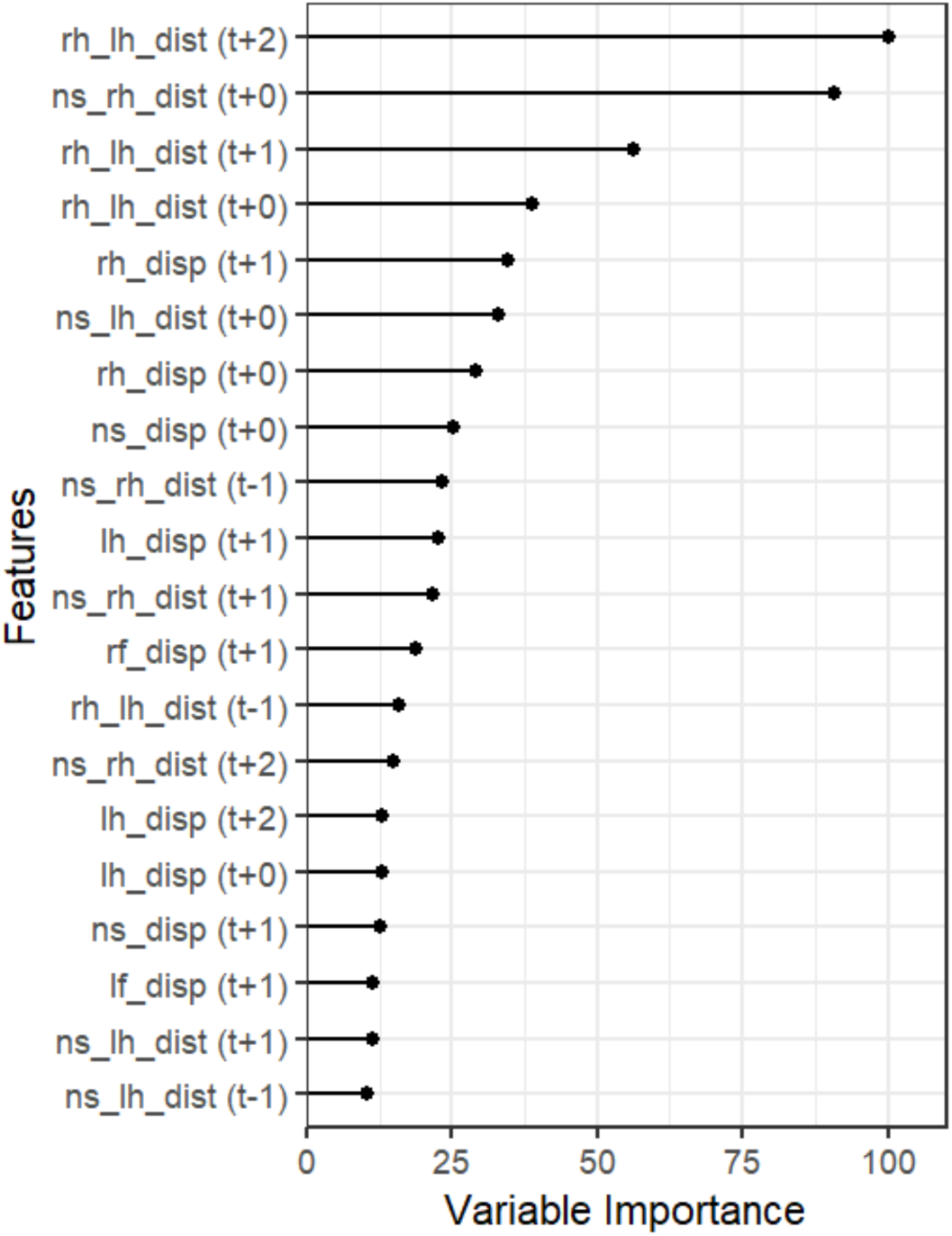
The importance of each feature in the final model. The top 20 features with the highest importance are selected and displayed. The values shown are relative values, with the most important of the entire set of features being 100. Finally, the temporal relationship of the feature used for each frame to the frame itself is shown as t + x or t − x (i.e., when x = 2, the feature was obtained from the two frames after the frame). ns = nose, rh = right front paw, lh = left front paw, rf = right rear pow, lf = left rear pow, disp = displacement, dist = distance.

In the test set, face washes (sensitivity = 0.69, positive predictive value (PPV) = 0.85) (Fig. 7A), and forelimb flails (sensitivity = 0.56 and PPV = 0.74) (Fig. 7B) were detected frame by frame with reasonable accuracy. In contrast, chin rubs (sensitivity = 0.16, PPV = 0.58) (Fig. 7C), gapes (sensitivity = 0), and head shakes (sensitivity = 0) were almost undetectable (Table 3). Others (i.e., movements other than the behaviors of interest) were detected with 0.99 sensitivity and 0.97 PPV. This prediction result was similar to that of the cross-validated prediction in the training set (Table 4).

**Figure 7.**
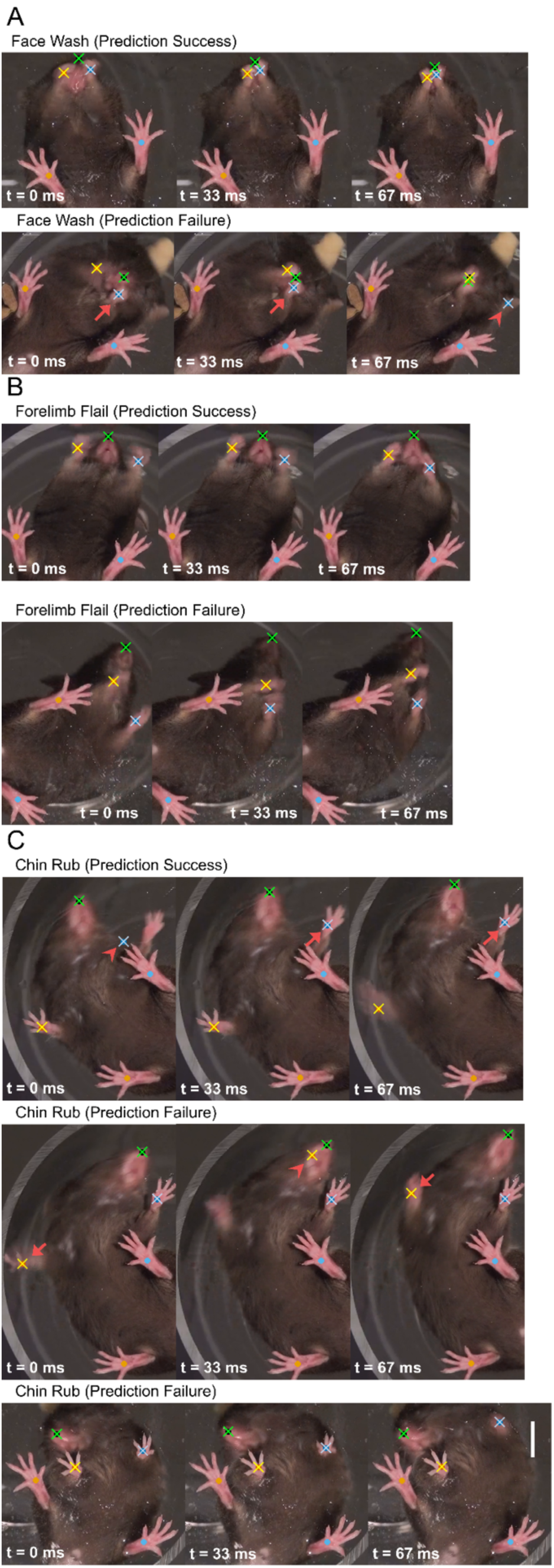
Typical examples of positive and false classifications by the present random forest classifier. The nose, right front paw, and left front paw of the mouse are highlighted with green, yellow, and blue crosses, while the right and left rear paws are marked with yellow and blue dots, respectively. Successive images were recorded at intervals of 33 milliseconds (ms) (1/30th of a second). Scale bar, 1 cm. **(A)** Examples of the classifier’s positive and false classifications of face wash. (Top) A series of frames in which the classifier correctly classifies the face wash. (Bottom) In this series, the classifier did not detect face wash, and there was a pose estimation error: at t = 0 and 33 ms, the right front paw positions were correctly estimated (red arrow), but at t = 67 ms, they were incorrectly estimated (red arrowhead). **(B)** Examples of the classifier’s positive and false classifications of forelimb flails. (Top) A series of frames in which the classifier correctly classified the forelimb flails. (Bottom) A series of frames in which the classifier did not detect forelimb flails and there was no pose estimation error. **(C)** Examples of the classifier’s positive and false classifications of chin rubs. (Top) A series of frames in which the classifier correctly classifies chin rubs. In this series, the left front paw of the mouse was moving so quickly that it could not be accurately recorded on the video; at t = 33 and 67 ms, its position was correctly estimated (red arrow), but at t = 0 ms, its position was incorrectly estimated (red arrowhead). However, pose estimation was successful. (Middle) In this series, the classifier did not detect chin rub, and there was a pose estimation error: at t = 0 and 67 ms, the right front paw positions were correctly estimated (red arrow), but at t = 33 ms, they were incorrectly estimated (red arrowhead). (Bottom) The classifier did not detect chin rubs in this series, and there was no pose estimation error.

**Table 3.** Observation and prediction of the number of frames that include each type of disgust reaction and other movement in the test set.

|  | Observation |  |  |  |  |  |  |
| --- | --- | --- | --- | --- | --- | --- | --- |
| Prediction |  | Chin<br>Rub | Face<br>Wash | Forelimb<br>Flail | Gape | Head<br>Shake | Others |
|  | Chin Rub | 45 | 0 | 0 | 0 | 0 | 33 |
|  | Face Wash | 0 | 283 | 25 | 0 | 0 | 26 |
|  | Forelimb<br>Flail | 0 | 49 | 496 | 0 | 1 | 121 |
|  | Gape | 0 | 0 | 0 | 0 | 0 | 0 |
|  | Head Shake | 0 | 0 | 0 | 0 | 0 | 0 |
|  | Others | 236 | 77 | 362 | 22 | 18 | 27006 |

**Table 4.** Observation and prediction of the percentual average counts overall cross validation that include each type of disgust reaction and other movement in the train set.

|  | Observation |  |  |  |  |  |  |
| --- | --- | --- | --- | --- | --- | --- | --- |
| Prediction |  | Chin<br>Rub | Face<br>Wash | Forelimb<br>Flail | Gape | Head<br>Shake | Others |
|  | Chin Rub | 0.2 | 0.0 | 0.0 | 0.0 | 0.0 | 0.1 |
|  | Face Wash | 0.0 | 1.4 | 0.2 | 0.0 | 0.0 | 0.2 |
|  | Forelimb<br>Flail | 0.0 | 0.2 | 2.7 | 0.0 | 0.0 | 0.6 |
|  | Gape | 0.0 | 0.0 | 0.0 | 0.1 | 0.0 | 0.0 |
|  | Head Shake | 0.0 | 0.0 | 0.0 | 0.0 | 0.0 | 0.0 |
|  | Others | 0.9 | 0.3 | 2.2 | 0.2 | 0.1 | 90.8 |

### Disgust reaction scoring

To evaluate the performance of the classifier for the disgust reaction score, predictions of the total number of frames that constitute individual disgust reactions were scored according to certain rules, as mentioned above, and the scored values were compared with the values manually counted and scored by a human. We found a high correlation between the scores calculated from the classifier’s prediction and human count (Pearson’s r = 0.98, Fig. 8, see Supplementary Movie).

**Figure 8.**
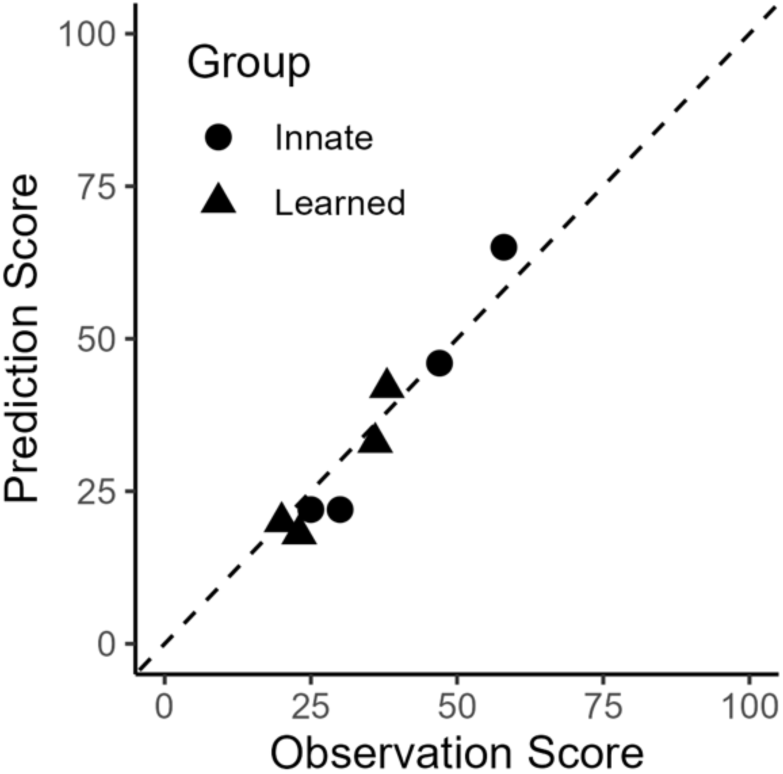
The classifier’s predictions were highly correlated with human observations. Comparison of disgust reaction scores for each video between observations and predictions in the test dataset. Pearson’s r = 0.98.

## Discussion

We constructed a classifier using DLC and random forest to automatically count innate and learned disgust reactions in the taste reactivity test. The total predicted values of the classifier showed a strong correlation with human observations. The present study is the first to automatically assess disgust reactions in taste reactivity tests.

Our classifier was constructed according to the published taste reactivity test protocols. Therefore, the scores analyzed by our classifier can be compared with manually counted scores. We found a very high correlation in this comparison (r = 0.98, Fig. 5). The accuracy of our classifier is comparable to that of previous automated methods: the pup retrieval behaviors in mice were automatically assessed with 0.51-0.90 correlations using DeepLabCut and machine learning [27], and the number of chemical-induced scratching in mice was predicted with 0.98 correlations using a convolutional recurrent neural network [29].

The present study used DLC for marker-less pose estimation because a previous study demonstrated excellent performance [30]. Nonetheless, alternative open-source software programs such as DeepPoseKit [31] and SLEAP [32] allow for comparable estimations. Our present classifier may also assess disgust reactions even if these programs are used instead of DLC. The research field of markerless pose estimation has made remarkable progress in recent years, and more accurate programs are likely to become available. Accordingly, automatic behavior evaluation procedures that perform pose estimation in advance, such as our present classifier, are expected to improve in accuracy because at least some of the failures of our classifier were a result of pose estimation failures (Fig. 6).

A recent study evaluated facial expressions in response to taste stimuli in mice using a machine learning-based method [33]. Facial expressions in response to bitter stimuli were automatically distinguishable from those in response to the other stimuli. Bitter-induced facial expressions were also induced by the intake of saccharin solution paired with intraperitoneal LiCl injection and by the intake of NaCl solution at high concentrations. However, it remains unclear what specific brain functions are reflected in the observed facial expressions because the taste stimuli used may evoke multiple brain functions, including disgust and motivation to avoid the stimulants. In addition, the facial expressions detected by this method in mice were challenging to interpret as relevant facial expressions in other species, including humans, making it difficult to interpret the mouse’s facial expressions as a readout of emotion. In contrast, all disgust reactions expressed by mice are shared by several different species, such as rats, monkeys, and human neonates [15]. This fact supports the idea that a mouse’s disgust reactions and human disgust may share a common neural basis. In support of this, taste reactivity may reflect different brain functions and neural activities from facial expressions because the numbers of disgust reactions and facial expressions do not always fit in mice [34]. In addition to facial expression, intake and licking behaviors have been widely examined in taste avoidance and evaluation studies, but the results from these behaviors do not always match those of taste reactivity [20, 35]. Thus, taste reactivity will continue to be a valuable readout for future research to understand the neural basis of disgust.

Our classifier can be built with minimal resources. In the video of the taste reactivity used in the present study, three of the five types of behavioral reactions that make up the disgust reaction, chin rub, face wash, and forelimb flail, accounted for 93.46% of the total disgust reactions (Fig. 3B). As these three reactions are characterized by the movement of the nose and the front and rear paws, tracking the five body parts of the mice, the nose, and the front and rear paws (Fig. 4A) seemed to be sufficient to track these reactions, and thus the disgust reactions observed in the present study. These five tracking points are generally smaller than those used in other studies using DeepLabCut (8-13 points/mouse) [28,30].

Although the behaviors detected in other studies are different from those in the present study and cannot be directly compared, the relatively small number of tracking points in the present study achieved automated tracking with a relatively small burden of manual annotation and machine training costs. In addition, an inexpensive and commercially available handheld video camera with low frame rates (30 frames per second), which is equal to or less than that used in previous studies (30-60 frames/sec) [29,36], was sufficient to record mouse behavior for subsequent analysis using our classifier. Using videos with low frame rates saves time for pose estimation and reduces the storage size of the original videos, ultimately improving the model [37].

However, our classifier could hardly detect gapes and head shakes, which are the other two reactions that constitute the disgust reaction (Tables 3 and 4). This may be because the tracking point for the gapes was not set at a sufficiently high level to detect movement around the oral cavity, which was assumed necessary for detection. For the head shake, the 30 fps record may have been insufficient because the movements were too fast. Although the inability to detect these reactions was not problematic in the present study because the frequency of these reactions was low in mice (Fig. 3), increasing the number of tracking points may be required to predict disgust reactions in rats, where gapes are more frequently observed [38].

In the present study, we developed an automated assessment method for mouse disgust reactions. The present method allows for the evaluation of disgust reactions independent of the skill level and bias of the human evaluator. In addition, the present method will facilitate the implementation of taste reactivity tests in large-scale screening and long-term experiments that require counting numerous disgust reactions, which are challenging to perform manually. Thus, the present method may help researchers implement taste reactivity tests in a broader range of studies than previously thought [20]. The present method is expected to accelerate our understanding of the neural basis of disgust, contribute to understanding the pathophysiology of various psychiatric disorders, and advance the development of preventive and therapeutic interventions.

## Acknowledgments

This work was supported by JST SPRING, Grant Number JPMJSP2120 to S.I., JSPS KAKENHI Grant Number JP19K06938, Takeda Science Foundation, TOBE MAKI Scholarship Foundation, and a TMDU Priority Research Areas grant to D.H.T. We thank Dr. Tsutomu Tanabe at TMDU for his helpful comments and discussion.

## Author contributions

N.U. supervised this project. S.I. and D.H.T. conceived the project and designed experimental and analytical strategies. D.H.T. conducted experiments. S.I. analyzed the data and wrote the code. S.I. drafted the manuscript, which was discussed and edited by all the co-authors.

## Data availability

The videos, code, and generated datasets used in the present study can be provided by the corresponding authors upon reasonable request.

## Additional Information

Competing Interests

The authors declare that they have no conflicts of interest.

