## Supplementary Movie Legend for "Automatic quantification of disgust reactions in mice using machine learning"

**Typical example of video recording in a mouse expressing disgust reactions.**

The nose, right front paw, and left front paw of the mouse are highlighted with green, yellow, and blue crosses, while the right and left rear paws are marked with yellow and blue dots, respectively. The left area shows the total number of disgust reactions and each type of disgust reaction, with the upper half counted by human observers and the lower half automatically determined by the classifier. A video was recorded at 30 fps.
